# Transcriptionally Active HIV Reservoirs Enriched for Interferon-Inducible APOBEC3-Related Mutational Signatures Associate with Arterial Inflammation

**DOI:** 10.64898/2026.06.03.729898

**Authors:** Kazuo Suzuki, Angelique Levert, Emma Yoo, Shady Abohashem, John Zaunders, Lucette A. Cysique, Hirotaka Ode, Yasumasa Iwatani, Ahmed Tawakol, Priscilla Y. Hsue, Felicia C. Chow, Bruce J. Brew

**Affiliations:** St Vincent’s Centre for Applied Medical Research, NSW State Reference laboratory for HIV, Sydney; St Vincent’s Clinical School, Faculty of Medicine, UNSW, Sydney, Australia; Cardiovascular Imaging Research Center, Massachusetts General Hospital and Harvard Medical School, Boston, USA; UNSW Psychology, Sydney, NSW, Australia; Departments of Neurology and Immunology and Peter Duncan Neurosciences Unit, St Vincent’s Hospital, University of New South Wales and University of Notre Dame Sydney; Clinical Research Center, National Hospital Organization Nagoya Medical Center, Nagoya, Aichi, Japan; Department of Microbiology and Immunology, Hamamatsu University School of Medicine; Department of Medicine (Cardiology), Massachusetts General Hospital and Harvard Medical School, Boston, USA; Department of Medicine (Cardiology), University of California, Los Angeles, Los Angeles, USA; Weill Institute for Neurosciences and Departments of Neurology and Medicine (Infectious Diseases), University of California, San Francisco, USA

**Keywords:** Cell-associated HIV-1 transcripts, Viral reservoir, APOBEC3, Cardiovascular Disease, Arterial Inflammation

## Abstract

Despite suppressive antiretroviral therapy (ART), people with HIV (PWH) remain at increased risk of cardiovascular disease (CVD), likely driven, at least in part, by persistent immune activation and vascular inflammation. Mechanistic pathways may include ongoing low-level HIV transcription within immune cells, particularly peripheral blood mononuclear cells (PBMCs), which may contribute to chronic inflammatory signaling. Persistent viral transcription and inflammatory signaling may also be associated with an interferon-responsive cellular environment conducive to APOBEC3 activation and mutational editing of viral transcripts. APOBEC3 enzymes, which are interferon (IFN)-inducible cytidine deaminases, generate characteristic premature stop codons (PSCs) within retroviral genomes and may reflect interferon-associated inflammatory states relevant to vascular pathology.

We investigated whether APOBEC3-associated PSC signatures and reservoir-associated drug resistance mutations (RA-DRMs) in viral reservoirs, as markers of persistent reservoir transcriptional activity, are associated with vascular inflammation in ART-suppressed individuals. PBMCs from ART-suppressed PWH (n=36) were analyzed for long HIV-1 *gag/pol* transcripts (>4.2 kb) and PSCs as a surrogate marker of APOBEC3 activity, with RA-DRMs assessed by Oxford Nanopore sequencing. Arterial inflammation was quantified using PET imaging of the aorta and carotid arteries. Additionally, an exploratory panel of >60 cardiometabolic, clinical, and inflammatory biomarkers was profiled to identify host immune signatures associated with reservoir PSC burden, with focused analyses on circulating inflammatory biomarkers sCD14, sCD163, and IP-10.

Long HIV-1 transcripts were detected in 67% (24/36) despite ART. Higher PSC frequency within these transcripts was significantly associated with increased aortic inflammation, but not carotid inflammation, in univariable analysis (β = 0.00834, p = 0.013). RA-DRMs were detected in 50% (12/24), including triple-class resistance, but were not associated with vascular inflammation or soluble biomarkers. In multivariable models, PSCs remained independently associated with aortic inflammation after adjustment for sCD14 and sCD163 (β = 0.0075, p = 0.023), and remained the only significant predictor when IP-10 was included (β = 0.0073, p = 0.036). These findings suggest that APOBEC3-associated PSC signatures in long HIV transcripts identify transcriptionally active HIV reservoir states associated with aortic inflammation during suppressive ART. Rather than reflecting ongoing mutational accumulation or direct pathogenic effects of PSCs themselves, these signatures may serve as biomarkers of an interferon-associated inflammatory reservoir state related to cardiovascular risk.

**Graphical Abstract:** 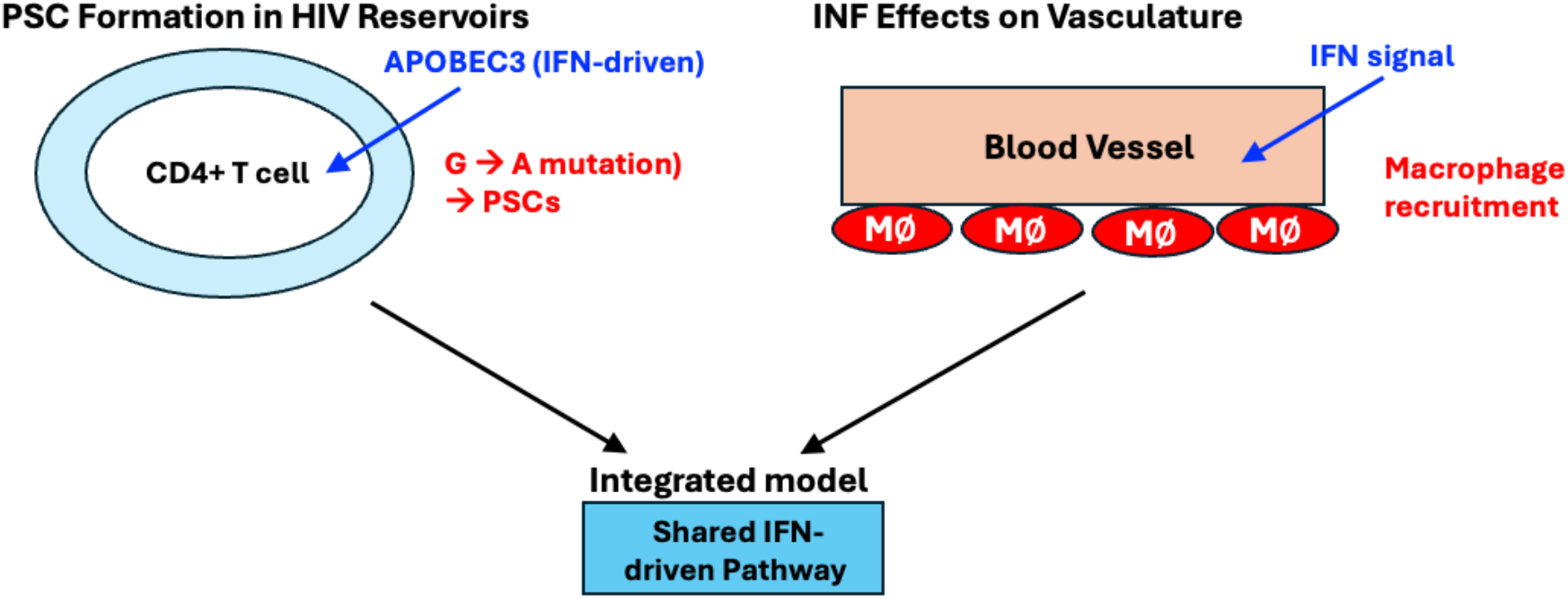

**Highlights: Interferon-inducible APOBEC3-associated PSC signatures in HIV reservoirs and vascular inflammation:**

- APOBEC3-associated premature stop codon (PSC) signatures were identified in transcriptionally active HIV reservoir-derived RNA despite suppressive ART.
- Higher PSC burden was significantly associated with increased aortic vascular inflammation measured by FDG-PET.
- Reservoir-associated drug resistance mutations (RA-DRMs) showed no association with arterial inflammation.
- PSC burden remained independently associated with aortic vascular inflammation in multivariable analyses.
- PSC-enriched reservoir states were associated with inflammatory biomarker patterns consistent with monocyte–macrophage activation and vascular immune activation.
- These findings are consistent with an interferon-inducible APOBEC3-edited reservoir state associated with persistent arterial inflammation during suppressive ART.
- APOBEC3-associated mutational signatures may represent a candidate biomarker of vascular inflammation and cardiovascular risk in people with HIV.

## Introduction

Despite effective suppressive antiretroviral therapy (ART), people with HIV (PWH) remain at increased risk of cardiovascular disease (CVD), including atherosclerosis and vascular inflammation ^1–3^. Importantly, this increased risk persists even in individuals with fully suppressed plasma viral load ^4–6^, indicating that mechanisms beyond active systemic viral replication contribute to vascular pathology ^7–9^.

Chronic immune activation and persistent inflammation are thought to play central roles in this process ^10^. Circulating markers of monocyte and macrophage activation, including soluble CD14 (sCD14), soluble CD163 (sCD163), and IP-10, have been associated with cardiovascular risk and arterial inflammation in prior studies ^5,10–15^. However, most of these studies include heterogeneous populations with variable viral suppression, and it remains unclear whether these associations persist in individuals with complete and durable viral suppression, where plasma HIV RNA is below the level of detection ^5,10,11,13–16^.

This distinction is important because in fully suppressed individuals, as in the absence of detectable viraemia, it remains uncertain whether these biomarkers may reflect residual immune dysregulation rather than ongoing viral replication, and their relationship to tissue-level vascular inflammation is not well defined. Therefore, understanding cardiovascular risk in this setting requires exploration of mechanisms that operate independently of detectable viremia.

Subclinical vascular inflammation in people with HIV has been demonstrated using ^18^F-fluorodeoxyglucose positron emission tomography (FDG-PET), including increased arterial inflammation in ART-treated individuals and associations with systemic inflammatory and neurocognitive outcomes ^5,11,12,17,18^. These findings are supported by a broader body of imaging studies, which consistently demonstrate persistent carotid and aortic inflammation in virologically suppressed PWH and indicate that vascular inflammation is associated with atherosclerotic disease burden ^1,5,12,17,18^. Importantly, these studies establish that vascular inflammation persists despite long-term ART-mediated viral suppression.

One potential mechanism underlying persistent CVD risk in people with PWH is ongoing low-level HIV transcription within long-lived cellular reservoirs, despite effective ART. Although ART suppresses plasma viraemia, it does not directly inhibit proviral transcription, and previous studies, including ours have shown that HIV transcripts commonly remain elevated in ART-suppressed individuals ^19–25^. While these transcriptional events may not produce infectious virus, persistent viral RNA expression may contribute to chronic innate immune activation through persistent viral RNA expression.

In this context, host antiviral restriction pathways active during the establishment of HIV reservoirs, particularly APOBEC3-mediated editing, may leave stable mutational signatures within integrated proviruses. APOBEC3 enzymes are interferon-inducible cytidine deaminases that generate characteristic G-to-A hypermutations, including premature stop codons (PSCs), during reverse transcription ^26–31^. Mutation signatures of these PSCs provide a molecular footprint of host antiviral activity within viral reservoirs ^32,33^ and may reflect long-lived virus–host interactions detectable during suppressive ART.

Although APOBEC3-mediated restriction has been implicated in viral evolution and host antiviral defence, whether APOBEC3-associated mutational signatures within transcriptionally active HIV reservoirs are associated with vascular inflammation in fully ART-suppressed individuals remains unknown. Similarly, the clinical relevance of reservoir-associated drug resistance mutations (RA-DRMs) persisting within transcriptionally active reservoirs in relation to cardiovascular outcomes has not been defined.

In this study, we specifically examined a cohort of individuals with fully suppressed plasma viral load to assess whether APOBEC3-associated PSC signatures and RA-DRM patterns within cell-associated HIV-1 (CA-HIV) transcripts relate to vascular inflammation measured by FDG-PET. We further evaluated a comprehensive panel of cardiometabolic and inflammatory biomarkers. This design provided an opportunity to investigate potential associations between reservoir-derived viral signatures and vascular inflammation in the absence of detectable plasma viremia.

## Results

### Detection of CA HIV-1 short transcripts in PBMCs

The demographics of the 36 individuals included in this study are presented in Table S1.

Participants had a mean age of 59.7 years and were predominantly white men (97%) with chronic, treated HIV-1 infection. All participants were on suppressive ART (median duration 33 years) with HIV-1 RNA below the limit of detection. The dataset of more than 60 biomarkers spanning cardiometabolic risk, clinical comorbidities, inflammatory pathways, including monocyte activation markers (sCD14 and sCD163), and the IFN-inducible chemokine IP-10 (CXCL10), reflecting IFN-driven immune activation is available via the link provided below (see method). The baseline CD4/CD8 ratio indicated partial immune reconstitution under ART (mean 1.04 ± 0.70), however substantial inter-individual variability was observed, ranging from profound immune dysregulation to near-complete normalization. This heterogeneity was consistent with persistent immune dysregulation in a subset of individuals despite effective ART.

Total RNA was extracted from stored PBMCs, followed by analysis of CA-HIV-1 short RNA transcripts. Short HIV-1 RNA transcripts, reflecting HIV-1 promoter (LTR) activity and enabling assessment of transcriptional activation in virally suppressed individuals, were detected in all 36 participants, with a median level of 8,817 copies per 106 PBMCs (Figure 1A, B). In contrast, plasma HIV-1 RNA was undetectable in all participants (Figure 1A). These findings are consistent with our previous reports ^23–25,34–36^, which demonstrated abundant short HIV-1 RNA transcripts within viral reservoir cells, indicating persistent transcriptional activity despite long-term suppressive ART.

**Figure 1.**
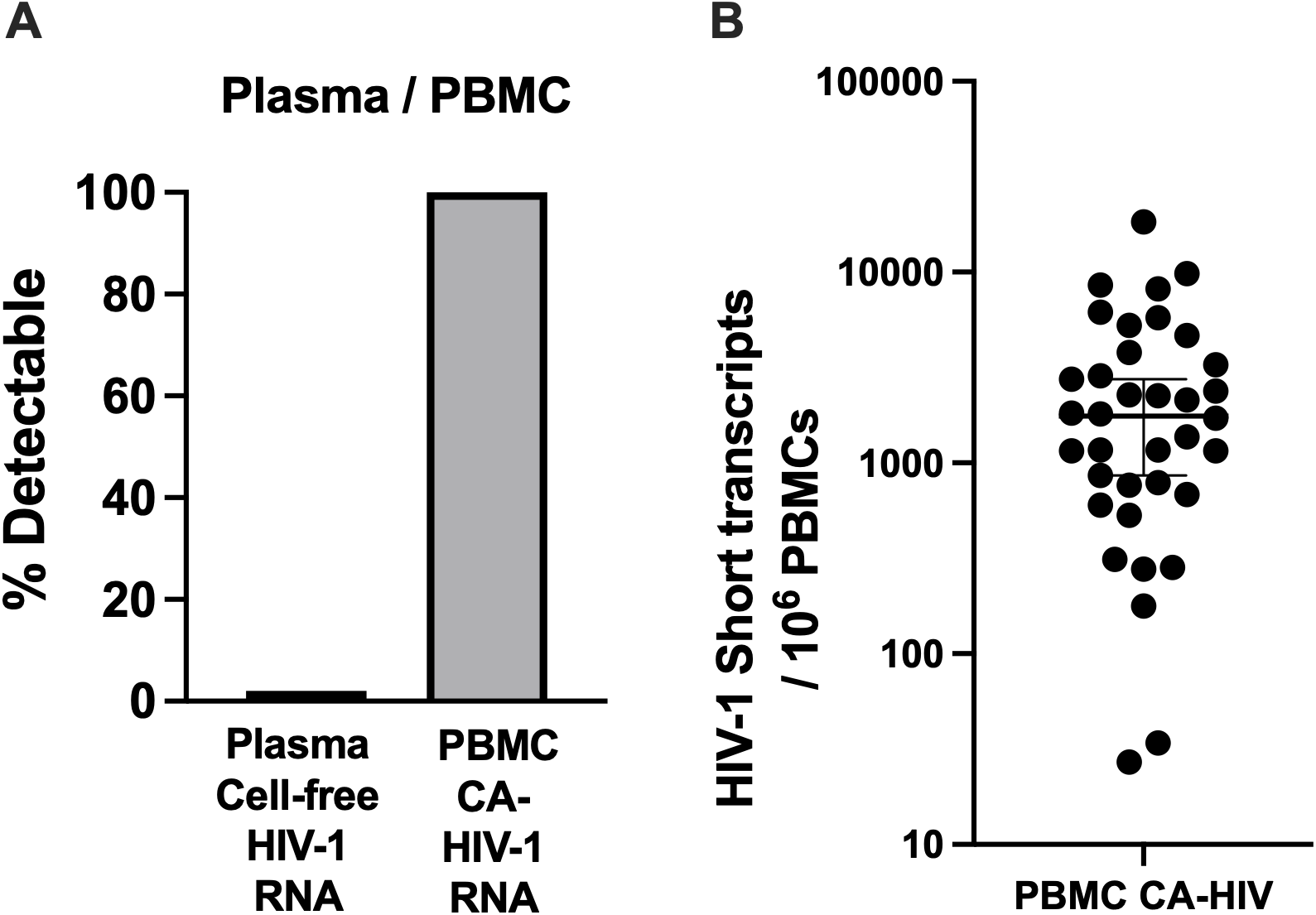
Detection of short HIV-1 RNA transcripts in PBMCs from people with HIV and suppressed plasma viral load. **(A)** Comparison of plasma HIV-1 RNA and CA-short HIV-1 RNA transcript levels. Plasma HIV-1 RNA was undetectable in all participants, whereas short HIV-1 RNA transcripts were consistently detected in all 36 individuals, indicating ongoing viral transcription despite suppressive antiretroviral therapy (ART). **(B)** Short HIV-1 RNA transcript levels in PBMCs, quantified as copies per 106 PBMCs (median, 8,817; 95% confidence interval [CI]).

We next examined associations between short HIV-1 RNA transcript levels and a comprehensive panel of biomarkers related to cardiometabolic risk, clinical comorbidities, inflammatory pathways, and monocyte activation. No significant associations were identified between short HIV-1 RNA transcript levels and these systemic biomarker measures.

### Validation of Nanopore HIV Drug Resistance Analysis Using Long *gag/pol* Sequences

Recent advances in Oxford Nanopore sequencing, particularly duplex-based approaches, have substantially improved accuracy (>99.9%) and demonstrated high concordance with Sanger-based HIV-1 DRM testing, supporting its application for viral genotyping ^37–41^. Nanopore sequencing also enables long-read analysis of viral haplotypes and within-host subpopulations, including low-frequency variants that may be missed by short-read approaches ^37–45^.

In this study, prior to application in the CVD cohort analyses, we performed independent analytical validation of the Nanopore workflow to ensure accurate detection of DRMs in both plasma-derived virus and cell-associated viral reservoirs in peripheral blood mononuclear cells for our validation of nanopore HIV DRM analysis.

We established a single-step RT-PCR assay targeting the HIV-1 *gag/pol* region (>4.2 kb). HIV-1 RNA was extracted from plasma and amplified using this single-step RT-PCR approach, as described in the Methods section. In the initial evaluation, we assessed amplification specificity and sensitivity using 26 plasma samples with viral loads ranging from 261 copies/mL to >1,000,000 copies/mL. The assay demonstrated robust and highly specific amplification performance following plasma RNA extraction (Figure S1A). Importantly, this sensitivity is below the conventional clinical threshold for HIV DRM testing (>2,000 copies/mL), indicating suitability for low-level viremia.

To validate DRM detection, we analysed 59 plasma samples with viral loads ranging from 2,240 to 3,150,000 copies/mL. Nanopore sequencing was used to identify DRMs and compared with the hospital’s standard method, which uses two-step RT-PCR targeting separate proteinase/reverse transcriptase (PR/RT) and integrase (IN) regions ^23^. Concordance between Nanopore sequencing and Sanger sequencing was ≥95% across all mutation classes, with complete agreement for major and accessory mutations in both PR/RT and integrase regions (Figure S1B). A representative comparison between Sanger and Nanopore sequencing is shown in Figure S1C. As expected, minor variants (<20% frequency) were detected by Nanopore sequencing but not by Sanger sequencing, highlighting the higher sensitivity of next-generation sequencing approaches. The full dataset is available via the provided link below. These results confirm that the Nanopore-based method achieves high accuracy and strong concordance with Sanger sequencing, consistent with previous reports using plasma samples ^40,43^.

We next extended the analysis to CA-HIV-1 long *gag/pol* transcripts from peripheral blood mononuclear cells of individuals with low-level viremia (LLV; 261 copies/mL; ID2137, Figure S1A) and those with undetectable plasma viral load (pVL; n=13). All CA HIV long *gag/pol* analyses were performed using nested PCR (see Methods).

In the LLV participant (ID2137), plasma analysis identified a single reverse transcriptase-region DRM, whereas CA-RNA analysis revealed major DRMs spanning three antiretroviral drug classes (Figure S1D). Among individuals with undetectable plasma viral load, long HIV RNA transcripts were detected in 46% (6/13). Of these, 83% (5/6) harboured at least one DRM affecting one or more drug classes, suggesting persistence of resistance-associated viral sequences despite plasma viral suppression (Figure S1D). These findings indicate the presence of reservoir-associated HIV-1 DRMs within cellular compartments during suppressive ART, supporting further investigation of their potential clinical relevance in the context of cardiovascular diseases.

### Persistent drug resistance mutations detected in long *gag/pol* transcripts

We applied a novel analytic approach to the CVD cohort to detect CA HIV-1 long *gag/pol* frame transcripts encoding “Reservoir-associated HIV-1 DRMs”. The long transcripts were successfully amplified in 24 of 36 individuals (67%) (Figure 2). Subsequently, we performed nanopore-based NGS to assess the presence of RA-DRMs and PSCs within the pol region of the 24 samples with detectable long transcripts (Figure 2). Full-length duplex reads were analyzed using a 5% variant frequency cutoff to capture potential low-frequency RA-DRMs in the CA viral mRNA (see Methods). PSCs were identified at variable frequencies across samples, and some samples contained pol sequences bearing RA-DRMs but no PSCs. Detected RA-DRMs within the putatively intact sequences were categorized into four classes: proteinase inhibitors (PIs), non-nucleoside reverse transcriptase inhibitors (NNRTIs), nucleoside reverse transcriptase inhibitors (NRTIs), and integrase strand transfer inhibitors (INSTIs) with prevalence illustrated using color-coded bars.

**Figure 2.**
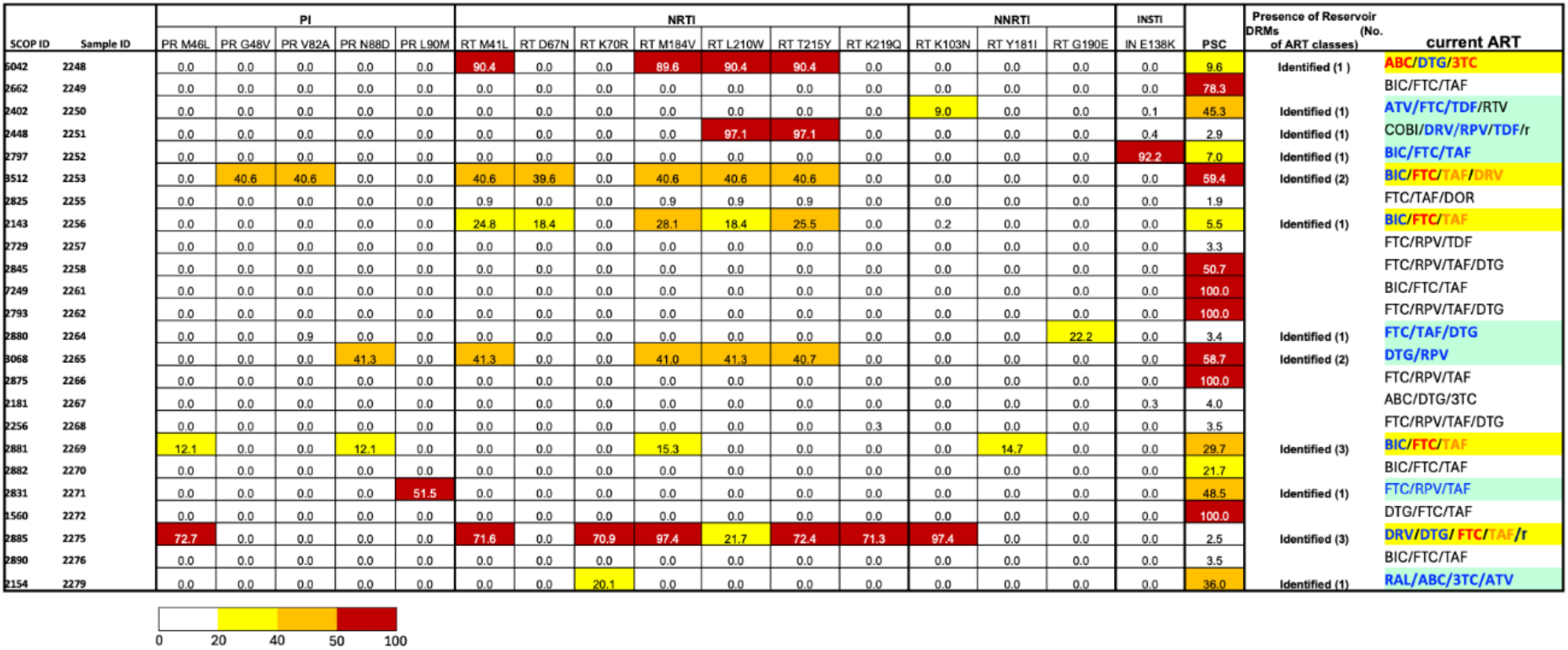
Persistent RA-DRMs Reflect Current ART Use. CA-HIV-1 long *gag/pol* RNA transcripts were successfully amplified in 67% (24/36) of participants and analyzed for RA-DRMs. RA-DRMs were detected in 50% (12/24) of individuals with amplifiable transcripts. These 12 individuals with detectable reservoir-associated RA-DRMs are shown by antiretroviral drug class (PI, NNRTI, NRTI) at the time of analysis. Identified RA-DRMs are aligned with current ART regimens. For each sample, the within-sample prevalence of major RA-DRMs in the pol region is shown, excluding sequences containing PSCs. The percentage shown for each sample represents the proportion of RA-DRMs consistent with selection by the current ART regimen.

Reservoir-associated resistance mutations were identified in 12 of 24 individuals and, when classified into mutually exclusive categories, comprised single-class resistance in 8 cases (NNRTI, n = 2; NRTI, n = 4; PI, n=1; INSTI, n=1), dual-class resistance in 2 cases (PI/NRTI, n = 2), and triple-class resistance in 2 cases (PI/NNRTI/NRTI, n=2). A single integrase substitution (E138K) in INSTI was detected in one individual (ID2252); however, this occurred in the absence of accompanying major INSTI resistance mutations. This isolated E138K substitution, is consistent with a potential APOBEC-associated G-to-A mutational process and, in this context, is unlikely to have functional significance for INSTI susceptibility.

Regarding the relationship between RA-DRMs and current antiretroviral therapy (ART), five individuals harboured RA-DRMs corresponding to components of their current ART regimen. These cases are highlighted in yellow in Figure 2, with ART agents linked to identified RA-DRMs shown in red and orange lettering, and estimated key regimen components indicated in blue lettering. In contrast, seven individuals exhibited RA-DRMs that did not correspond to their current ART regimen (highlighted in blue in Figure 2), generally at low variant frequencies in next-generation sequencing (NGS) data (<25%), with the exception of the integrase substitution E138K in ID2252 (92.2%) and the PI mutation L90M in ID2271 (51.5%).

Importantly, a high frequency of RA-DRMs exhibiting thymidine analogue mutation (TAM) patterns associated with prior zidovudine (AZT) and lamivudine (3TC) exposure was identified within the NRTI resistance profiles. These TAM-associated resistance patterns were observed in 4 of the 5 individuals highlighted in yellow (RA-DRMs corresponding to components of the current ART regimen) and in 2 of the 7 individuals highlighted in blue (RA-DRMs not corresponding to the current ART regimen) (Figure 2). These findings may support long-term persistence of archived NRTI resistance signatures.

We next examined associations between identified RA-DRMs and a comprehensive panel of biomarkers related to cardiometabolic risk, clinical comorbidities, inflammatory pathways, and monocyte activation. No statistically significant associations were observed, suggesting that the detected reservoir resistance patterns were not directly linked to systemic inflammatory or metabolic signatures in this cohort. Given the lack of clinical correlates, we subsequently investigated viral sequence integrity in long HIV-1 RNA transcripts, focusing on PSCs. This analysis was performed to explore whether defective viral genomes and host restriction-associated mutational processes, including potential APOBEC-associated G-to-A hypermutation, contributed to the observed reservoir architecture.

### Premature stop codons in the HIV reservoirs

CA-HIV-1 long *gag/pol* transcript analysis using Oxford Nanopore sequencing enabled identification of PSCs within individual long reads as described above. We next examined associations between PSC burden and a comprehensive panel of biomarkers, including arterial inflammation. Of 36 individuals assessed, 12 were excluded from PSC analysis due to insufficient abundance of long HIV-1 RNA transcripts required for reliable assessment of sequence integrity. The inability to detect long HIV-1 RNA transcripts in this subset likely reflects extremely low reservoir transcriptional output rather than absence of proviral HIV DNA sequences. Accordingly, PSC analyses were performed in 24 individuals with detectable long-read HIV-1 RNA transcripts.

### Association between APOBEC3-associated PSCs and arterial inflammation

An exploratory analysis of >60 cardiometabolic, clinical, and inflammatory biomarkers was first performed to comprehensively assess host factors potentially associated with reservoir-derived viral signatures. Following this global screening approach, markers of monocyte activation (sCD14 and sCD163) and the IFN-inducible chemokine IP-10 were selected for focused secondary analyses based on their established relevance to HIV-associated immune activation and vascular inflammation ^5,10–15^. These host biomarker profiles were subsequently evaluated in relation to imaging-based measures of arterial inflammation..

We next examined the association between APOBEC3-associated PSC signatures in long HIV transcripts and arterial inflammation measured by FDG-PET. PSC burden was significantly associated with increased aortic inflammation, as assessed by ^18^F-fluorodeoxyglucose uptake (p = 0.013; linear regression analysis; Figure 3A), whereas no significant association was observed between PSCs and carotid TBRmax (Figure 3B).

**Figure 3.**
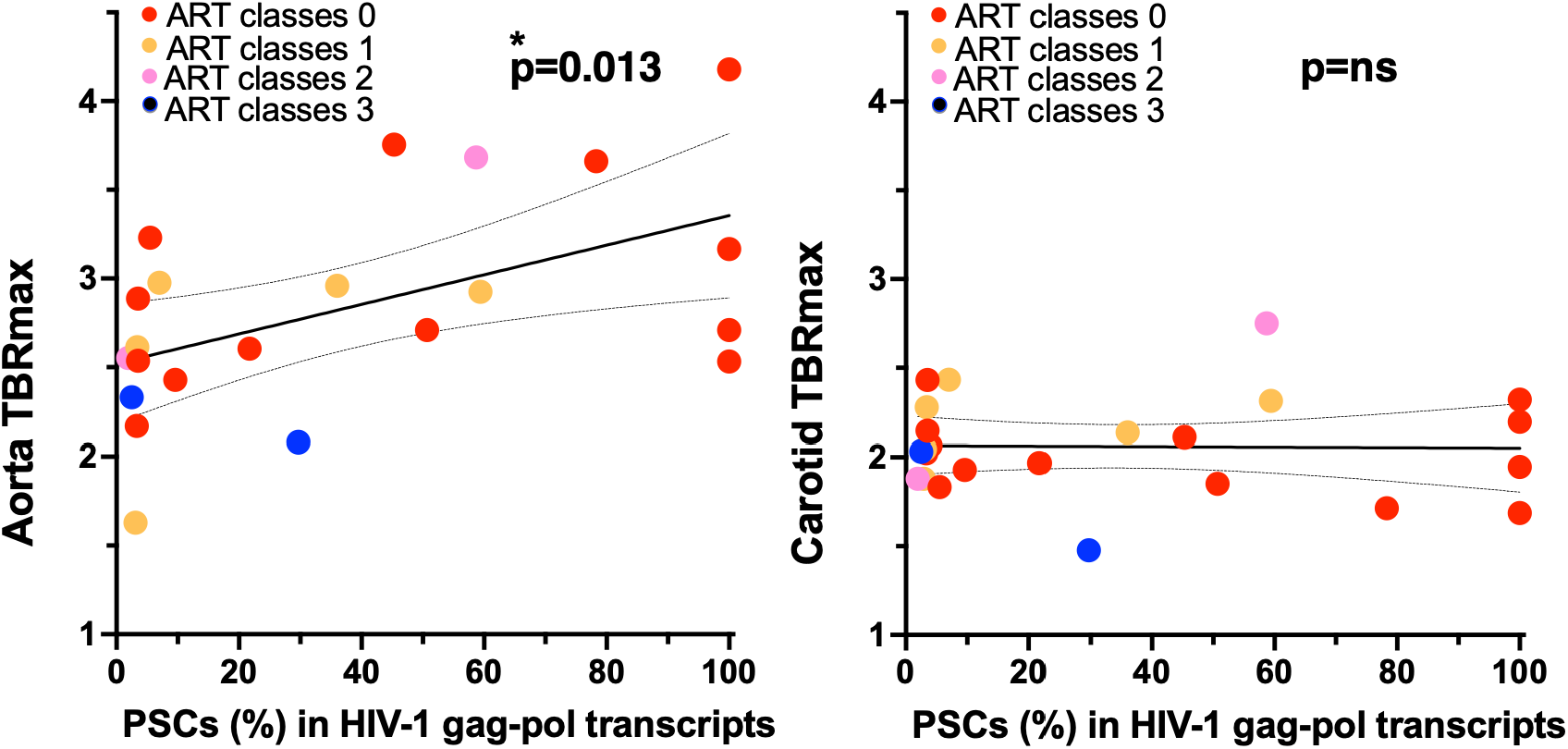
Association between APOBEC3-associated PSCs and arterial inflammation. Aortic (**A**) and carotid (**B**) arterial inflammation were assessed using ^18^F-FDG-PET imaging and expressed as TBRmax (maximum target-to-background ratio), defined as arterial wall ^18^F-FDG uptake normalized to blood pool activity within the region of interest. Higher TBRmax values indicate greater arterial inflammation. Identified RA-DRMs within viral reservoirs are shown across antiretroviral therapy (ART) classes. Note: Aortic TBRmax values were not available for two individuals (ID2251 and ID2267).

In comparison, the number of ART classes represented by RA-DRMs showed no association with aortic TBRmax, supporting a selective association between APOBEC3-associated PSC signatures and aortic inflammation.

### Univariable association analysis

We next performed univariable linear regression analyses to assess associations between PSC burden and markers of monocyte activation (sCD14 and sCD163), as shown in Table 1. PSC burden demonstrated a significant positive association with the outcome (β = 0.0083, 95% CI 0.0020–0.0147; p = 0.013). In contrast, no significant associations were observed for sCD14 (β = 0.0005, 95% CI −0.0012 to 0.0007; p = 0.605) or sCD163 (β = −0.0119, 95% CI −0.260 to 0.283; p = 0.928). Overall, PSC burden was the only variable showing a statistically significant association in univariable analysis.

**Table 1.** Univariable Association Analysis.

| Variable | $\beta$ | 95% CI | p-value | Sig | Interpretation |
| --- | --- | --- | --- | --- | --- |
| **PSCs** | 0.0083 | 0.0020 – 0.0147 | **0.013** | **Yes** | **Significant positive association** |
| **sCD14** | 0.0005 | –0.0012 – 0.0007 | 0.605 | No | No significant association |
| **sCD163** | –0.0119 | –0.260 – 0.283 | 0.928 | No | No significant association |

We then constructed a parsimonious adjusted model defined a priori, incorporating age and smoking status as covariates (Table 2). In this model, PSC burden remained independently associated with the outcome (β = 0.0075, 95% CI 0.0013–0.0137; p = 0.020), whereas neither age (β = −0.0204, 95% CI −0.0511 to 0.0103; p = 0.18) nor smoking status (β = −0.8583, 95% CI −1.963 to 0.2465; p = 0.12) showed significant associations. These findings indicate that the association between PSC burden and arterial inflammation remained robust after adjustment for key demographic and behavioral covariates.

**Table 2.**
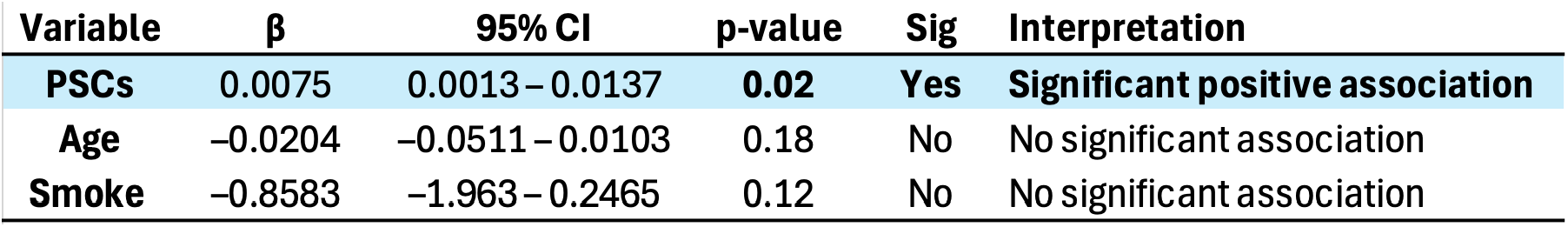
Parsimonious Adjusted Model (a priori: Age and Smoking)

| Variable | $\beta$ | 95% CI | p-value | Sig | Interpretation |
| --- | --- | --- | --- | --- | --- |
| **PSCs** | 0.0075 | 0.0013 – 0.0137 | **0.02** | **Yes** | **Significant positive association** |
| **Age** | –0.0204 | –0.0511 – 0.0103 | 0.18 | No | No significant association |
| **Smoke** | –0.8583 | –1.963 – 0.2465 | 0.12 | No | No significant association |

### Multivariable association analysis

We subsequently performed multivariable linear regression analyses incorporating PSC burden together with monocyte activation markers (sCD14 and sCD163), as shown in Table 3. PSC burden remained significantly associated with the outcome in the multivariable model (β = 0.0075, 95% CI 0.0012–0.0139; p = 0.023), indicating an independent positive association. sCD14 was also independently associated with the outcome (β = 4.011, 95% CI 0.1085–7.914; p = 0.045), whereas sCD163 was not significantly associated (β = −0.0036, 95% CI −0.0157 to 0.0085; p = 0.539).

**Table 3.** Multivariable Model Including Inflammatory Markers.

| Variable | $\beta$ | 95% CI | p-value | Sig | Interpretation |
| --- | --- | --- | --- | --- | --- |
| **PSCs** | 0.0075 | 0.0012 – 0.0139 | **0.023** | **Yes** | **Significant positive association** |
| **sCD14** | 4.011 | 0.1085 – 7.914 | 0.045 | **Yes** | **Significant positive association** |
| **sCD163** | –0.0036 | –0.0157 – 0.0085 | 0.539 | No | No significant association |

Overall, PSC burden and sCD14 demonstrated independent positive associations with arterial inflammation, whereas sCD163 showed no significant effect.

We further extended the multivariable model to include the IFN-inducible chemokine IP-10 (Table 4). In this expanded model, PSC burden remained significantly associated with the outcome (β = 0.0073, 95% CI 0.00056–0.0142; p = 0.036). In contrast, no significant associations were observed for sCD14 (β = 3.388, 95% CI −0.854 to 7.63; p = 0.108), sCD163 (β = 0.001, 95% CI −0.0139 to 0.0159; p = 0.886), IP-10 (β = −0.00236, 95% CI −0.00656 to 0.00184; p = 0.247), or age (β = −0.00359, 95% CI −0.0429 to 0.0347; p = 0.847). These findings indicate that the association between PSC burden and arterial inflammation remained robust after adjustment for additional inflammatory and demographic variables.

**Table 4.** Further Multivariable models including inflammatory markers.

| Variable | $\beta$ | 95% CI | p-value | Sig | Significance |
| --- | --- | --- | --- | --- | --- |
| **PSCs** | 0.0073 | 0.00056 – 0.0142 | 0.036 | **Yes** | **Significant positive association** |
| **sCD14** | 3.388 | –0.854 – 7.63 | 0.108 | No | Not significant |
| **sCD163** | 0.001 | –0.0139 – 0.0159 | 0.886 | No | Not significant |
| **IP10** | –0.00236 | –0.00656 – 0.00184 | 0.247 | No | Not significant |
| **Age** | –0.00359 | –0.0429 – 0.0347 | 0.847 | No | Not significant |

## Discussion

In this study, we demonstrate that APOBEC3-associated PSC signatures within CA-HIV-1 transcripts from PBMCs are independently associated with aortic inflammation in people with HIV on suppressive antiretroviral therapy. This association persisted across multivariable models adjusting for established markers of monocyte/macrophage activation, including sCD14, sCD163, and IP-10 ^5,10–15^, suggesting that PSC burden reflects a biological signal not fully captured by systemic inflammatory biomarkers. In contrast, classical inflammatory mediators showed limited or inconsistent independent associations, highlighting the relative specificity of transcriptionally active reservoir clones containing APOBEC3-associated mutational signatures in relation to vascular inflammation.

Importantly, our study specifically focused on individuals with fully suppressed plasma viral load. In this setting, residual inflammatory markers may reflect chronic immune dysregulation rather than ongoing productive viral replication, and their relationship to tissue-level vascular inflammation remains incompletely defined. Understanding cardiovascular risk in this population therefore requires investigation of biological processes operating independently of detectable viraemia. Our findings suggest that reservoir-associated PSC signatures may provide insight into such processes beyond those captured by conventional plasma biomarkers.

Our results further support the concept that low-level HIV transcription may persist within long-lived cellular reservoirs despite successful ART. While these transcriptional events may not generate replication-competent virus, they may nevertheless contribute to chronic innate immune activation through persistent viral RNA expression. The persistence of PSC signatures as the only consistent predictor across multivariable models suggests that APOBEC3-associated mutational signatures may identify reservoir states associated with vascular inflammation that are not fully reflected by conventional systemic inflammatory biomarkers. In this context, PSCs may capture a more specific reservoir-derived signal than circulating inflammatory biomarkers, which likely reflect broader systemic immune activation.

Biologically, APOBEC3-mediated editing occurs during reverse transcription prior to proviral integration, generating characteristic G-to-A hypermutations and premature stop codons that become fixed within integrated viral genomes ^26–31^. Accordingly, PSC signatures detected in long HIV transcripts are unlikely to represent ongoing mutational accumulation or direct editing of residual transcripts. Rather, they likely reflect transcriptional activity of long-lived reservoir clones carrying archived APOBEC3-edited proviruses. The observed association between PSC burden and aortic inflammation therefore suggests that transcriptionally active APOBEC3-edited reservoirs may reflect a persistent IFN-associated inflammatory milieu linked to cardiovascular risk, rather than a direct pathogenic effect of PSCs themselves.

RA-DRM analysis provides additional insight into the biological nature of these reservoir-derived transcripts. In a subset of cases, resistance mutations corresponding to current ART regimens were not detected, suggesting that the observed transcripts are unlikely to reflect ongoing treatment-driven viral evolution. Instead, thymidine analogue mutation (TAM) patterns associated with historical zidovudine (AZT) and lamivudine (3TC) exposure support the persistence of archived viral genomes within long-lived reservoir cells. When considered alongside PSC signatures, these findings suggest that the viral reservoir contains a mosaic of historically selected and host-edited proviral genomes that remain transcriptionally active despite durable plasma viral suppression.

Importantly, similar TAM patterns linked to historical AZT and 3TC exposure were also identified in a companion parallel study examining neuroaxonal integrity associated with transcriptionally active HIV reservoirs, which has also been submitted to bioRxiv (Suzuki et al., submitted), further supporting the long-term persistence of archived, transcriptionally active reservoir populations across distinct HIV-associated comorbidities.

The selective association with aortic, but not carotid, inflammation suggests potential regional heterogeneity in vascular susceptibility to immune-mediated injury. Importantly, carotid inflammation was not significantly elevated in this cohort, which may explain the absence of an observable relationship between PSC burden and carotid TBRmax. Distinct inflammatory pathways and immune cell composition have been reported across vascular beds, supporting the concept that inflammatory responses may differ along the arterial tree ^46^. The aorta may be particularly prone to monocyte–macrophage recruitment and activation, as resident macrophages within the arterial intima, together with recruited mononuclear phagocytes, play central roles in the initiation and progression of aortic inflammation and atherosclerosis ^47^, while FDG-PET vascular uptake largely reflects macrophage metabolic activity within inflamed arterial tissue ^48^.

More broadly, these findings are consistent with emerging concepts from autoimmune and chronic inflammatory diseases ^49–52^, where persistent innate immune activation in the absence of ongoing infection is increasingly recognized as a contributor to endothelial dysfunction, arterial inflammation, and accelerated cardiovascular disease. In conditions such as Rheumatoid Arthritis, Systemic Lupus Erythematosus, and Psoriasis, sustained INF-associated immune activation has been linked to vascular injury and increased cardiovascular risk despite effective control of primary disease activity. Our findings raise the possibility that transcriptionally active, APOBEC3-edited HIV reservoir clones may similarly identify a persistent inflammatory reservoir state that parallels mechanisms of tissue-specific vascular injury observed in non-infectious inflammatory disorders.

Several limitations should be considered. The cross-sectional design precludes inference of causality, and the sample size limits power for detection of modest associations, particularly in expanded multivariable models. Additionally, while PSC signatures provide a useful molecular marker of APOBEC3-edited reservoir clones, they do not directly quantify APOBEC3 enzymatic activity or define the cellular source of transcription. Longitudinal studies incorporating tissue-based analyses will be required to clarify the temporal relationship between reservoir PSC burden and vascular inflammation.

In summary, APOBEC3-associated PSC signatures detected within HIV reservoirs are independently associated with aortic inflammation in treated HIV infection. These findings suggest that transcriptionally active APOBEC3-edited reservoir clones may reflect a persistent inflammatory reservoir state associated with cardiovascular risk, highlighting PSC signatures as a potential biomarker of vascular inflammation in people with HIV despite complete plasma viral suppression.

## Materials and Methods

### Participants

Thirty-six individuals with HIV (Table S1) were enrolled from a neurologic sub-study of a clinical trial evaluating the impact of alirocumab on cardiovascular risk. Inclusion criteria for the parent trial included: age ≥40 years, undetectable plasma viral load on ART, and at moderate to high cardiovascular risk defined as a history of CVD [e.g., prior myocardial infarction (MI) or stroke, coronary heart disease, peripheral arterial disease] or of at least one cardiometabolic comorbidity (e.g., hypertension, hyperlipidemia, diabetes mellitus, cigarette smoking). The study was approved by the UCSF Institutional Review Board, and written informed consent was obtained from all study participants. Participants with stored PBMCs close to the date of FDG-PET were eligible for this study.

To investigate host biological factors potentially associated with HIV reservoir activity, we analyzed a panel of more than 60 biomarkers spanning cardiometabolic risk, clinical comorbidities, and inflammatory and immune activation pathways. Particular attention was given to biomarkers relevant to monocyte activation (sCD14 and sCD163) and interferon-responsive immune signaling, including the interferon-inducible chemokine IP-10 (CXCL10). The complete biomarker dataset is available through the repository described below.

### ^18^F-FDG PET imaging and vascular inflammation quantification

Participants underwent FDG-PET of the neck and chest the University of California, San Francisco Imaging Center at China Basin on a GE Discovery STE PET/computed tomography (CT) scanner. A low-dose CT scan was performed for attenuation correction before PET acquisition. FDG uptake corrected for background venous activity was assessed in the bilateral carotid arteries and ascending aorta using previously validated methods ^5,11,12,17,18^. Briefly, measurements were made in the axial plane, and maximum standardized uptake values (SUVmax) were recorded from pre-defined sections of the carotid and ascending aorta walls. The target-to-background ratio (TBRmax) was defined as the ratio of the mean of the SUVmax measurements along the length of the target vessel to the background venous activity derived from either the internal jugular veins (for correction of carotid values) or the superior vena cava (for correction of aortic values). Higher TBRmax values indicate greater arterial inflammation.

As part of an exploratory systems-level phenotyping approach, more than 60 biomarkers were assessed alongside PET imaging of the aorta and carotid arteries, spanning multiple domains including inflammatory cytokines and chemokines (IP-10, MCP-1, IL-6, hsCRP), monocyte/macrophage activation markers (sCD163, sCD14), mental health and psychosocial burden (Perceived Stress Scale [PSS], PTSD, ACEs, PHQ-9), functional status (activities of daily living [ADL_now, ADL_best]), neuropsychological performance (Hopkins Verbal Learning Test [HVLT], Digit Span, Trail Making Test A and B, Stroop test, Controlled Oral Word Association

Test [COWAT], pegboard performance, and global cognitive z-scores), HIV clinical and immunovirological parameters (CD4/CD8 ratio, plasma HIV RNA, CD4 count, ART regimen, year of HIV diagnosis), and traditional cardiovascular disease risk factors and therapies (BMI, myocardial infarction, coronary heart disease, peripheral artery disease, family history, diabetes, hyperlipidemia, hypertension, stroke, lipid profile [cholesterol, LDL, HDL], HbA1c, HCV antibody status, and use of aspirin, statins, clopidogrel, and antihypertensive therapy). The full dataset is available via the provided link below.

### HIV reservoir analysis and molecular sub-studies supporting CVD cohort analyses

Before initiating the CVD cohort analyses (n = 36), the Oxford Nanopore sequencing workflow was established and technically validated using independently stored PBMC samples from St Vincent’s Hospital. These samples were used solely for assay development and were not included in subsequent clinical association analyses. This component was approved by the St Vincent’s Hospital Human Research Ethics Committee (HREC/15/SVH/425).

In the 36 participants included in the CVD imaging cohort, HIV-1 molecular analyses included blinded quantification of short HIV RNA transcripts in peripheral blood mononuclear cells (PBMCs), consistent with the reservoir transcription analyses reported in the Results section.

Subsequent analyses expanded to evaluate within-host viral genetic variation, including DRMs and premature stop codons (PSCs) within reservoir-derived HIV sequences (see sections below).

### Exploratory sequencing and extended reservoir analyses

As part of the SVH molecular sub-studies conducted prior to the CVD cohort analyses, Exploratory virological investigations were performed to define DRMs in plasma samples (see sections below). These analyses included:

i. 26 plasma samples analyzed for near full-length HIV-1 *gag/pol* regions (>4.2 kb), and
ii. 59 plasma samples analyzed for DRMs using Oxford Nanopore long-read sequencing.

These exploratory datasets informed the downstream reservoir stratification analyses reported in the Results (DRMs, PSCs, and transcriptional activity).

The reservoir analysis was further extended to a subset of participants (n = 14), including individuals with low-level viraemia (LLV; 261 copies/mL; ID2137), enabling comparative analysis of transcriptional and sequence-level persistence across individuals with durable viral suppression (plasma viral load suppressed for >2 years without documented viral blips) and those with intermittent low-level viraemia (documented viral blips within the preceding two years). These samples were used solely for assay development and were not included in subsequent CVD cohort clinical association analyses. These studies were approved by the St Vincent’s Hospital Human Research Ethics Committee (HREC LNR/16/SVH/327).

### Sample source and clinical definitions for SVH-based sub-studies

All samples were derived from routine standard-of-care clinical collections submitted to the NSW State Reference Laboratory for HIV at St Vincent’s Hospital (2020–2025), for routine monitoring of plasma viral load and CD4 T-cell counts.

Viral blips were defined as transient plasma HIV-1 RNA elevations between 20 and 200 copies/mL during routine clinical monitoring prior to the index analysis timepoint (Figure S1D). This definition aligns with the clinical stratification used in the Results when comparing blip vs fully suppressed individuals.

### Isolation of peripheral blood mononuclear cells (PBMC)

PBMCs were isolated from anti-coagulated blood by density centrifugation using Ficoll-Paque Plus (GE Healthcare, Chicago, IL, USA). Isolated PBMCs were cryopreserved in heat-inactivated, filter-sterilized bovine serum containing 10% dimethyl sulfoxide (DMSO) and stored in the vapor phase of liquid nitrogen.

### Short HIV-1 RNA transcripts analysis

RNA extraction and quantification of short HIV-1 RNA transcripts were performed as previously described ^23–25,34–36^. Primers and probes targeted the highly conserved “R” region within both the 5′ and 3′ long terminal repeats (LTRs; Double-R assay), enabling detection of total spliced and unspliced transcripts by one-step reverse transcriptase PCR (RT-PCR; WO2018/045425, PCT/AU2017/050974) as a measure of HIV-1 promoter activity. Amplicons were detected using precision image analysis of πCode MicroDiscs (PlexBio) on the IntelliPlex platform, as described previously ^23,35^.

### Long HIV-1 RNA transcripts analysis

A long-range RT-PCR protocol was used for selective amplification of full-length, in-frame *gag/pol* transcripts (>4.5 kb), as outlined in Australian PCT Application: PCT/AU2025/051218. Briefly, long RNA transcripts were amplified using a one-step RT-PCR protocol followed by nested long-range DNA PCR. This approach generated >4.5 kb amplicons of the *gag/pol* frame region for downstream sequencing.

### Nanopore Sequencing analysis

DNA libraries for nanopore sequencing were prepared as previously reported ^43^. Briefly, RT-PCR amplicons from long *gag/pol* transcripts were purified using the AMPure XP beads (Beckman Coulter, Brea, CA, USA), and their concentration was measured using a Qubit 4 Fluorometer with the Qubit 1X dsDNA HS Assay Kit (Thermo Fisher Scientific). To prepare DNA library, the amplicons were end-repaired, and then ligated with native barcoded and adapters. Long Fragment Buffer was used for the final wash. The prepared DNA library was quantified using Qubit and then loaded onto an R10.4.1 flow cell (Oxford Nanopore Technologies (ONT)). Sequencing was performed on a MinION Mk1C device using MinKNOW software (ONT), with fast base-calling enabled for a one-hour run.

### Analyses of the Drug resistance mutations

Based on duplex reads obtained from nanopore sequencing for each sample, prevalence of DRMs and PSCs within pol were examined as previously described with a slight modification ^43^. In this study, the major DRMs listed in Stanford HIV database (https://hivdb.stanford.edu/) were adopted for the analyses. DRMs that could arise from APOBEC3 editing were defined as shown in the previous report ^43^. To explore the DRMs and PSCs from the reads, reads were mapped on a partial HXB2 sequence (GenBank ID: K03455, 2,253-5,096 nt) as the reference. Then, reads that fully cover sequences between the 1st codon of protease (PR) and the 264th codon of integrase (IN) (2,253-5,021 nt according to HXB2 coordinate) were selected, because all the major DRMs appear between the PR 30 codon and IN 263 codon. Coverage for selected reads exceeded 100 per sample, consistent with prior work ^43^. Analyses focused on the region between PR codon 1 and IN codon 264. For each sample, per-site prevalence of major DRMs and PSCs was determined.

Additionally, prevalence of genotypes harboring unique DRM patterns, with or without PSCs, was assessed. A 5% cutoff was applied to detect DRMs, justified as follows: (i) duplexed reads with bidirectional strand analysis; (ii) >100 reads per site, with at least 100× depth, so a 5% variant is supported by ≥5 reads; (iii) low-frequency variants (5–10%) may be clinically significant ^53^, especially in ART-experienced individuals ^54^.

### Statistical analysis

HIV-1 RNA transcriptional activity in peripheral blood mononuclear cells (PBMCs) was quantified as previously described ^35^. Standard curves were generated from serial dilutions of HIV-1 plasmid DNA using GraphPad Prism v10 (GraphPad Software, San Diego, CA, USA). Cell-associated (CA) HIV-1 short RNA transcripts were quantified using the Double-R assay and expressed as copies per 106 PBMCs. Continuous variables were assessed for normality prior to analysis. Normally distributed variables are presented as mean ± standard deviation, and non-normally distributed variables as median with interquartile range. Categorical variables are presented as counts and percentages.

Following quantification of vascular inflammation, associations between ^18^F-FDG PET– derived TBRmax (aortic and carotid) and HIV-related molecular and clinical variables were evaluated.

Continuous variables were assessed for normality using visual inspection of distribution plots. Normally distributed data are presented as mean ± standard deviation, and non-normally distributed data as median with interquartile range. Categorical variables are presented as counts and percentages.

### Univariable analysis

Univariable associations between vascular inflammation (aortic and carotid TBRmax) and HIV reservoir-related molecular markers (including CA-HIV-1 RNA transcripts, frequency of RA-DRMs, and PSCs) were assessed using Pearson correlation for normally distributed variables and Spearman rank correlation for non-normally distributed variables. Linear regression analyses were additionally performed to estimate effect sizes, reported as β coefficients with 95% confidence intervals (CIs).

### Multivariable analysis

Multivariable linear regression models were constructed to determine whether HIV reservoir-associated molecular features were independently associated with vascular inflammation. Models were adjusted for established cardiovascular risk factors, including age, sex, smoking status, body mass index (BMI), hypertension, diabetes mellitus, and lipid-lowering therapy, where applicable.

Standardized β coefficients, 95% CIs, and two-sided p values were reported for all multivariable models. Model assumptions, including linearity, homoscedasticity, and multicollinearity, were assessed using residual diagnostics and variance inflation factors.

### Additional analyses

Exploratory analyses included the association between aortic TBRmax and frequency of PSCs using Pearson correlation.

All statistical analyses were performed using GraphPad Prism v10 (GraphPad Software).

A two-sided p value <0.05 was considered statistically significant.

## Acknowledgments

We thank the participants for their time and involvement in the study. We gratefully acknowledge trial coordination by PBMC biobanking by Kate Merlin, Sara Evans, Maeve Hodgson, Kevin Tran, and Ethan Lewis.

## Database

Complete biomarker measurements and associated datasets will be made available through the repository described in the final published article.

## Author Contributions

K.S. conceived the project. K.S., F.C., and B.J.B. designed the experiments and analyses. A.L., E.Y., and K.S. conducted the experiments. A.L., E.Y., Y.I., and H.O. performed the nanopore data analysis and procedures for long transcript analysis using nanopore sequencing. S.A, A.T., and P.Y. provided essential supports in arterial inflammation. L.C. provided essential supports in neurocognitive markers. K.S., F.C., and B.J.B. wrote the manuscript. H.O. and Y.I. reviewed, analyzed, and generated all nanopore-based datasets. J.Z. reviewed the immunology relevance of the findings. H.O. reviewed and confirmed the molecular analysis. B.J.B. and F.C. reviewed and confirmed the clinical relevance of the findings. All authors reviewed and approved the final manuscript.

---

**Supplementary Figure 1.**
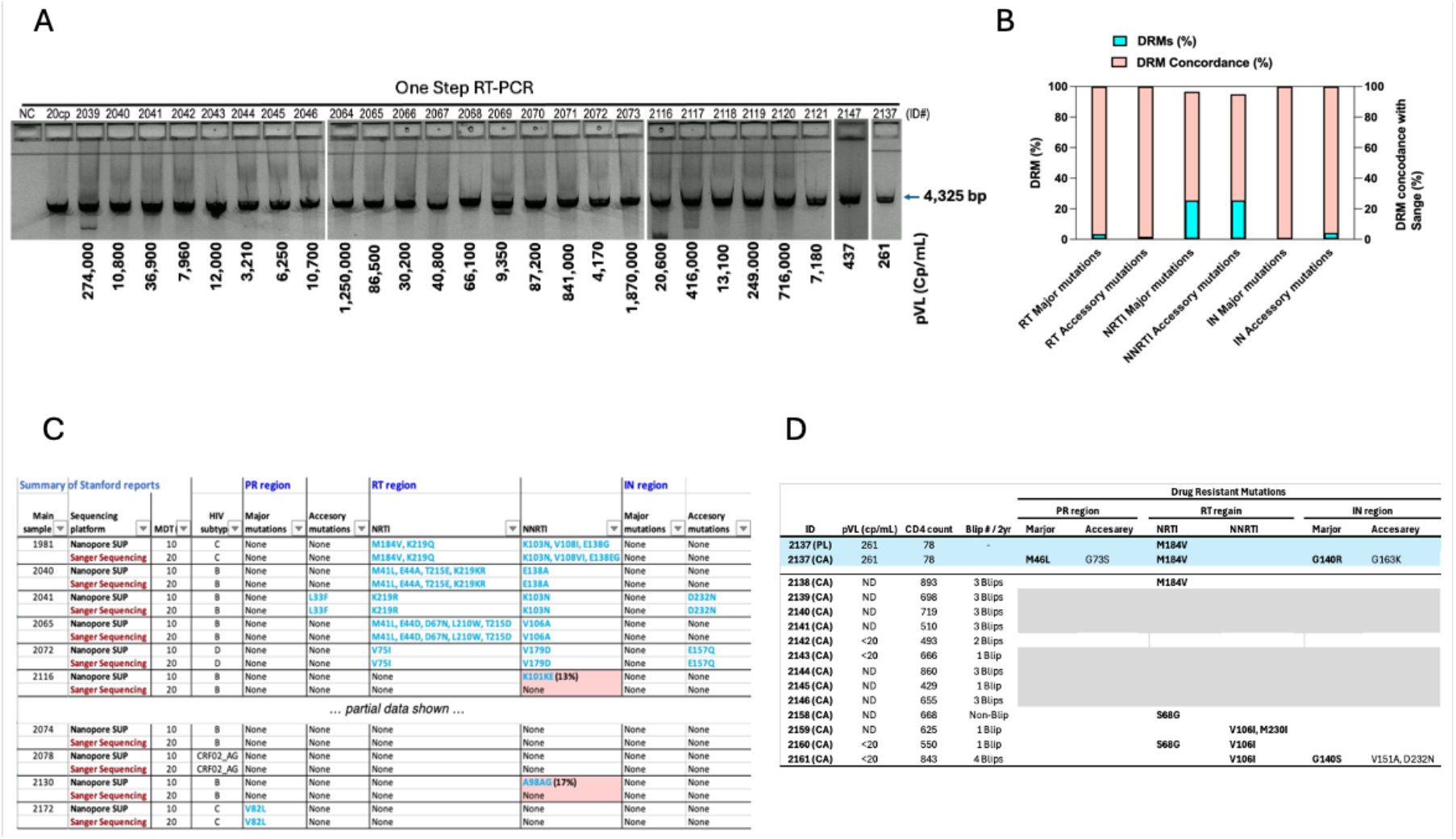
Analysis of long HIV-1 *gag/pol* RNA in plasma and in the reservoir-associated HIV-1 DRMs of 13 individuals. (**A**). Detection of long HIV-1 *gag/pol* RNA in plasma samples. Amplified PCR products were visualized by gel electrophoresis. (**B**). Frequency of DRMs across mutation classes (RT major, RT accessory, NRTI major, NNRTI accessory, IN major, IN accessory), and concordance (%) between Nanopore sequencing and Sanger sequencing. (**C**). Comparison of plasma sequencing results obtained by Sanger and Nanopore sequencing. Minority variants (<20% DRM frequency) were detected by Nanopore sequencing but not by Sanger sequencing, highlighting the increased sensitivity of long-read sequencing. (**D**). Detection of HIV DRMs in CA-RNA transcripts from viral reservoirs. One plasma sample with low-level viremia (261 copies/mL; ID2137) and thirteen samples with undetectable plasma viral load were analyzed. Long HIV RNA transcripts were detected in 46% (6/13) of individuals with suppressed viremia, of which 83% (5/6) harbored at least one DRM.

**Supplementary Table 1.**
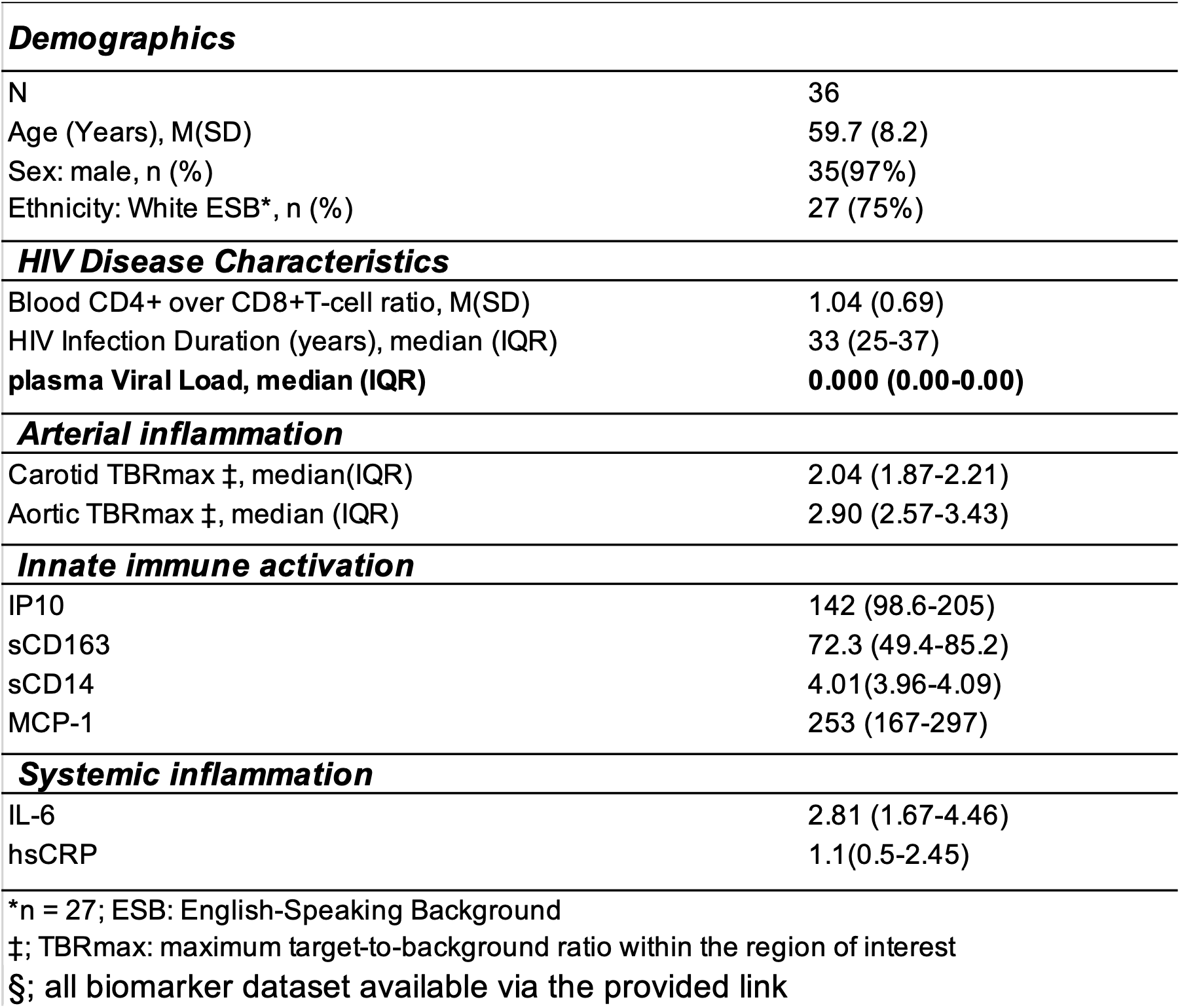
Cohort Demographics and Disease Characteristics.

|  |  |
| --- | --- |
| ***Demographics*** |  |
| N | 36 |
| Age (Years), M(SD) | 59.7 (8.2) |
| Sex: male, n (%) | 35(97%) |
| Ethnicity: White ESB*, n (%) | 27 (75%) |
| ***HIV Disease Characteristics*** |  |
| Blood CD4+ over CD8+T-cell ratio, M(SD) | 1.04 (0.69) |
| HIV Infection Duration (years), median (IQR) | 33 (25-37) |
| **plasma Viral Load, median (IQR)** | **0.000 (0.00-0.00)** |
| ***Arterial inflammation*** |  |
| Carotid TBRmax ‡, median(IQR) | 2.04 (1.87-2.21) |
| Aortic TBRmax ‡, median (IQR) | 2.90 (2.57-3.43) |
| ***Innate immune activation*** |  |
| IP10 | 142 (98.6-205) |
| sCD163 | 72.3 (49.4-85.2) |
| sCD14 | 4.01(3.96-4.09) |
| MCP-1 | 253 (167-297) |
| ***Systemic inflammation*** |  |
| IL-6 | 2.81 (1.67-4.46) |
| hsCRP | 1.1(0.5-2.45) |
| *n = 27; ESB: English-Speaking Background |  |
| ‡; TBRmax: maximum target-to-background ratio within the region of interest |  |
| §; all biomarker dataset available via the provided link |  |

